# homologizer: Phylogenetic phasing of gene copies into polyploid subgenomes

**DOI:** 10.1101/2020.10.22.351486

**Authors:** William A. Freyman, Matthew G. Johnson, Carl J. Rothfels

## Abstract

**Summary:**

1. Organisms such as allopolyploids and F1 hybrids contain multiple distinct subgenomes, each potentially with its own evolutionary history. These organisms present a challenge for multilocus phylogenetic inference and other analyses since it is not apparent which gene copies from different loci are from the same subgenome and thus share an evolutionary history.
2. Here we introduce `homologizer`, a flexible Bayesian approach that uses a phylogenetic framework to infer the phasing of gene copies across loci into their respective subgenomes.
3. Through the use of simulation tests we demonstrate that `homologizer` is robust to a wide range of factors, such as incomplete lineage sorting and the phylogenetic informativeness of loci. Furthermore, we establish the utility of `homologizer` on real data, by analyzing a multilocus dataset consisting of nine diploids and 19 tetraploids from the fern family Cystopteridaceae.
4. Finally, we describe how `homologizer` may potentially be used beyond its core phasing functionality to identify non-homologous sequences, such as hidden paralogs or contaminants.

## Introduction

Some individual organisms—such as allopolyploids and F1 hybrids—contain multiple distinct subgenomes, each with its own evolutionary history (in the case of polyploids, the homology of these subgenomes is related to a genome duplication event and they are referred to as “homeologous”). Such organisms present a particular challenge for multilocus phylogenetic inference because a researcher must take care to avoid assuming that gene copies from different loci share an evolutionary history, *i*.*e*., that they are from the same subgenome. For example, a diploid F1 hybrid individual will have one copy of a given nuclear locus from species A, and another from species B. If the copy of one locus from species A is treated as sharing an evolutionary history with the copy of another locus from species B (*e*.*g*., by mistakenly assigning them to the same haplotype in concatenated or multi-species coalescent analysis), the underlying bifurcating model will be violated and the inferences will be unreliable (McDade, 1990, 1992; Oxelman et al., 2017; Rothfels, 2021). This issue is not limited to multilocus phylogenies of single-copy nuclear loci—the same problem applies when attempting to include organellar and nuclear loci in a common analysis, or when analyzing ITS sequences with other loci.

Phasing copies across loci is related to the problem of haplotype assembly—the phasing of sequencing reads within a locus: in both cases, the goal is to avoid chimeric data that are a mix of multiple evolutionary histories. In the assembly problem, however, a researcher can rely on physical linkage to determine which reads belong to which haplotype (Kates et al., 2018; Schrinner et al., 2020; Majidian et al., 2020; Nauheimer et al., 2021; Tiley et al., 2021). This approach is not available in the locus-phasing case, where the loci are separated from each other by unsequenced regions; the only information available to determine whether two gene copies come from the same subgenome is in the phylogenetic history itself. Note that we use the term “gene copies” to refer both to copies that evolved in separate subgenomes (and thus do not share the same evolutionary history) and those that arose from allelic variation within the same subgenome (and thus do share the same species-level evolutionary history).

The need to phase loci is particularly prevalent in plants, since over one-third of extant species are inferred to be recent polyploids (Wood et al., 2009; Såstad, 2005). However, the issue is not restricted to plants. For example, both insects and diatoms have been reported to have an extensive history of polyploidy (Li et al., 2018; Parks et al., 2018), all salmonid fish are ancestrally polyploid (Alexandrou et al., 2013), and squamates and amphibians have clades with frequent allopolyploid lineage formation (Bogart, 1980; Bogart and Licht, 1986; Hedges et al., 1992; Lowe and Wright, 1966).

Historically, groups with extensive polyploidy have been under-studied phylogenetically (Soltis et al., 2014). Often, polyploids are dropped from phylogenetic analyses, in a “diploids-first” (or diploids-only) approach (*e*.*g*., Beck et al., 2010; Govindarajulu et al., 2011; Lee et al., 2002) or, if polyploids are included, authors tend to infer gene trees for each locus individually (*e*.*g*., Rothfels et al., 2014; Sousa et al., 2016; Chrtek et al., 2019; Melichárková et al., 2019; Griffin et al., 2011; Sessa et al., 2012a; Fortune et al., 2008; Rousseau-Gueutin et al., 2009; Kao et al., 2020). By omitting polyploids from multilocus analyses, we are not just removing the information those accessions could provide about general evolutionary patterns. Rather, we are systematically biasing our studies against groups with lots of polyploids, and we are impairing our ability to investigate questions where polyploidy may itself be an important factor (Ramsey and Ramsey, 2014; Mayrose et al., 2015; Rothfels, 2021). For example, if we are unable to include polyploids in multilocus phylogenies, we are limited in our ability to investigate classic questions in evolutionary biology such as the impact of polyploidy on diversification rates (Stebbins Jr, 1940; Wagner Jr., 1970; Soltis et al., 2009; Mayrose et al., 2011; Zhan et al., 2014; Tank et al., 2015; Freyman and Höhna, 2018; Zenil-Ferguson et al., 2019), or the association of polyploidy with niche breadth or with dispersal and establishment ability (Stebbins Jr, 1985; Ramsey and Schemske, 2002; Marchant et al., 2016).

In theory, researchers could get around this locus-phasing problem by inferring individual gene trees, and then using the phylogenetic position of the copies to determine to which subgenome they belong (the “phase” of the copies). For example, if a polyploid always has two gene copies, one of which is closely related to diploid species A, and the other to diploid species B, and this is true across all loci sampled, then one could confidently conclude that the “A” copies share one evolutionary history, and the “B” copies another (*e*.*g*., Sessa et al., 2012b; Dauphin et al., 2018). However, given variable amounts of missing data (such as failure to recover one of the copies for a subset of the loci, or failure to sequence the related diploids for all loci) and the phylogenetic uncertainty inherent in inferring singlegene phylogenies, this method can be exceedingly difficult to apply, especially in datasets with many polyploids, or with many loci (Bertrand et al., 2015).

Beyond this by-eye approach, and approaches that rely upon existing reference sequences, such as that of Hénocq et al. (2020) and Nauheimer et al. (2021), there are, to our knowledge, three currently available methods for phasing copies across loci. First, Bertrand et al. (2015) developed an approach to phase two loci by finding the largest set of sequence pairs such that any incongruence between the two resulting gene trees could be due to stochastic or coalescent error. A somewhat similar approach was developed and refined by Oberprieler and colleagues (Oberprieler et al., 2017; Lautenschlager et al., 2020). Instead of explicitly testing for sources of incongruence, this method looks for the subgenome assignments of copies within a locus, and then across loci, that minimizes the number of inferred deep coalescent events in the context of the phylogeny of the diploid species. Both methods are relatively fast, although the complexity of the analysis increases quickly with the number of polyploid accessions and with the number of loci. Perhaps more fundamentally, these methods add in accessions sequentially, so the phylogenetic information in the full dataset is not available to inform the phasing of each polyploid copy.

In contrast to the two methods above, which require an iterative series of analyses and are based on particular test statistics, the third available method, `alloPPnet` (Jones et al., 2013; Jones, 2017) is a one-step parametric Bayesian model for the inference of polyploid species trees (more accurately, species networks) under the multi-species coalescent. As such it explicitly models single polyploid formation events, takes into account all the data from the full sample, and, for example, requires that the post-polyploidization subgenome lineages have the same effective population size. The chief limitation of `alloPPnet` is that it cannot accommodate ploidy levels higher than tetraploid.

Here we introduce `homologizer`, a simple, flexible, tree-based method to infer the posterior probabilities of the phasing of gene copies across loci that can be performed on a fixed topology, or while simultaneously estimating the phylogeny under a wide range of phylogenetic models, including the multi-species coalescent. Users can thus apply `homologizer` to infer the phylogeny while treating the phasing as a nuisance parameter, to infer the phasing while integrating out the phylogeny, or for any combination of these goals. This method can utilize the full dataset of all loci jointly as opposed to phasing each locus individually (information about topology and subgenome identity from each locus is thus available to inform the phasing of the other loci), does not require any external data (such as a reference genome), and does not require that the progenitor diploids of the polyploid accessions be extant or sampled.

`homologizer` is implemented in the open-source Bayesian inference software `RevBayes` (Höhnaet al., 2016). It consists of dedicated MCMC proposals that switch tip assignments among gene copies and MCMC monitors that log the sampled phasing assignments. These `homologizer` features can be included in conjunction with any of the other modular phylogenetic inference components that `RevBayes` offers, enabling the phasing to be estimated under a large variety of models (*e*.*g*., models that assume a shared gene tree for some or all loci, multi-species coalescent models that do or do not assume a constant population size, etc.). We anticipate that `homologizer` will be of greatest interest to phylogeneticists, but its applications extend beyond phylogenetics to other cases where it is important to determine from which subgenome a particular gene copy originates, for example, studies of expression-level dominance and subfunctionalization (Edger et al., 2018).

## Methods

`homologizer` makes use of multilocus sequence data, where a locus is a homologous genomic region present across the accessions sampled for the analysis (though the method allows for missing data). The output of the method is the posterior distribution of phased homeologs, *i*.*e*., the posterior distribution of the assignments of each gene copy, for each locus, into each of the available subgenomes. If joint inference of the phasing and phylogeny is performed, the posterior distribution of the multi-locus phylogeny is also inferred, along with all other parameters of the model.

### The homologizer model

The `homologizer` model represents phylogenies containing polyploids as multilabeled trees (“mul-trees”), where each polyploid accession is present multiple times, once for each subgenome (each distinct evolutionary history; Huber et al., 2006). Gene copies are phased into subgenomes for each locus by swapping which gene copy is assigned to each of that accession’s tips in the mul-tree (Figure 1). For each polyploid accession, there are (*n*!)^*g*^ ways to phase the gene copies, where *n* is the number of gene copies per locus and *g* is the number of loci. The `homologizer` model thus adds a set of extra latent variables—gene-copy phase parameters—to the underlying phylogenetic model:

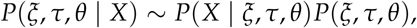

where *ξ* are the gene-copy phase parameters, *τ* is the tree, *θ* are all the other parameters of the model, and *X* is the set of sequence alignments.

**Figure 1:**
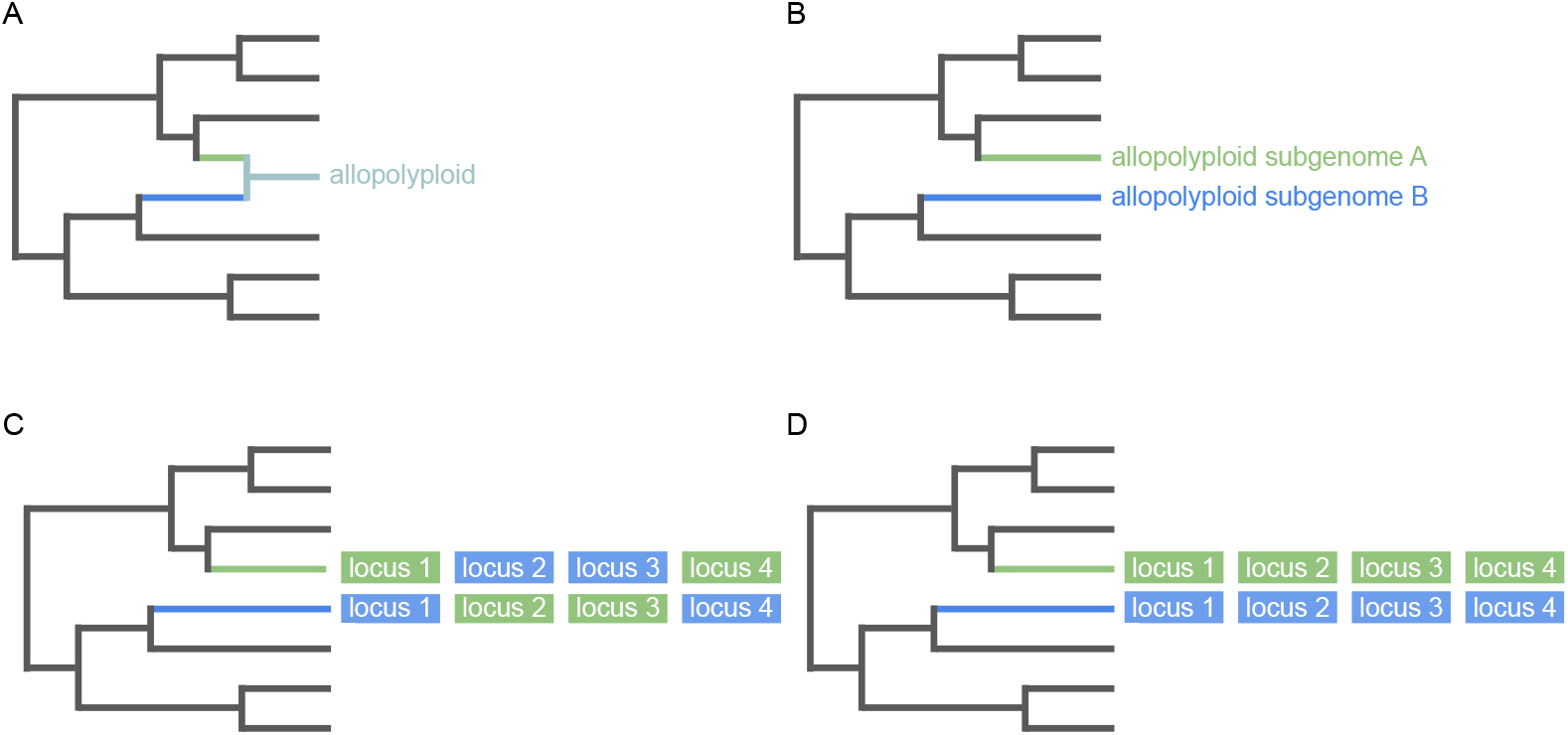
Phasing gene copies into polyploid subgenomes on a mul-tree. *A*: A phylogenetic network representing a single reticulation giving rise to an allopolyploid. *B*: The mul-tree representation of this phylogenetic network has two leaves (green and blue) representing the two subgenomes of the allopolyploid. *C*: Four loci were sequenced from the allopolyploid. Two copies (green and blue) of each locus were recovered. Loci 2 and 3 are incorrectly phased. *D*: After phasing, each locus is assigned to the correct subgenome.

The user defines the set of gene-copy phase parameters *ξ*, which for each accession and each locus consists of *n* 1 parameters for the *n* gene copies to be phased. Thus, the number of such parameters increases with both the number of polyploid subgenomes and with the number of loci. A given gene-copy phase parameter can take *n* values, corresponding to the *n* possible assignments of sequences to a given tip in the mul-tree; these values are each assigned an equal prior probability. The prior probability of the tree *P*(*τ*) is specified using a standard phylogenetic model such as a birth-death process or a uniform prior on the topology with exponential priors on branch lengths. The other parameters of the model, *θ*, represent a nucleotide substitution model. For example, one may specify the GTR substitution model (Tavaré, 1986) with exchangeability rates and stationary frequencies sampled from flat Dirichlet priors. The posterior probability *P*(*ξ, τ, θ* | *X*) is then estimated by MCMC (Metropolis et al., 1953).

### Summarizing homologizer inferences

The phasing estimates from a `homologizer` MCMC analysis can be summarized and plotted using the `R` (R Core Team, 2013) package `RevGadgets` (Tribble et al., 2020), which displays the joint maximum *a posteriori* (MAP) phasing assignment across the set of a polyploid’s subgenomes for each locus and summarised on a phylogeny. This joint MAP assignment is the highest probability assignment of each gene uniquely to a subgenome. Additionally, we show the marginal posterior probability for each locus and each subgenome. These marginal posterior probabilities are useful when quantifying the uncertainty in the joint MAP phasing assignment. For example, imagine a hexaploid with three subgenomes (A, B, and C) and three gene copies at a particular locus. If subgenomes A and B are difficult to distinguish (*e*.*g*., they could be sister to each other), then `homologizer` may infer a low posterior probability for the joint MAP phasing assignment of this locus. However, examining the marginal probabilities of each phasing assignment would reveal that subgenome C was phased with high posterior probability and that the low joint posterior probability was due to the difficulties in phasing between subgenomes A and B. Furthermore, the mean of the marginal probabilities across loci of the phasing assignment for a subgenome summarizes the model’s overall confidence in the phasing for that subgenome; this information is also incorporated into the `RevGadgets` visualization (one should also consider examining the median marginal probabilities for each subgenome). Note that, in contrast to the joint MAP phasing, the marginal MAP phasing for each individual subgenome may result in assignment of the same gene copy to multiple subgenomes.

### Comparing homologizer models

Users interested in distinguishing gene copies that evolved in separate polyploid subgenomes from those that arose from allelic variation within the same subgenome (or that are otherwise non-homeologous) can set up a series of `homologizer` models—which differ in the number of mul-tree tips available for phasing—and select the best fitting model using statistical model comparisons. To perform `homologizer` phasing model comparisons, the user can run a stepping-stone analysis (Xie et al., 2011) to compute the marginal likelihood of each model. The best fitting model is then selected using Bayes factors (Kass and Raftery, 1995).

Consider the example in Figure 2. Here, the polyploid accession has two subgenomes, was sequenced for four loci, and for each of those loci, two copies were recovered. It could be that each locus is represented by one copy from each of the subgenomes and thus the accession should be given two tips in the mul-tree (panel A in Figure 2). However, it’s possible, instead, that the copies at one or more of the loci are haploid allelic variants from one of the subgenomes. In this case, three mul-tips are needed—two tips for one subgenome, and one tip for the other subgenome (panel B in Figure 2). To compare these two models (one with two mul-tree tips for this accession, and one with three) using Bayes factors, the data must be kept constant, and only the number of phase-assignment parameters changed. This condition is satisfied by running both models with a “third set” of blank sequences added for this accession, and assigned to a mul-tree tip, but with that tip phased only in the “three-tip” analysis. In the two-tip analysis, conversely, the phase of this tip is not estimated and is fixed to blank sequences; after the analysis it can be pruned from the tree sample.

**Figure 2:**
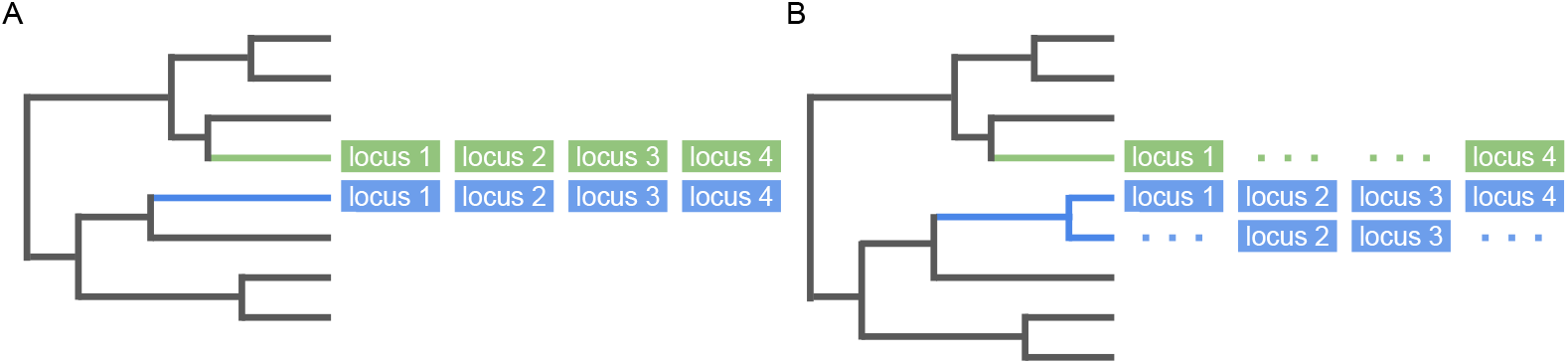
Model testing for allelic variants and/or homeologs. In this example, two copies of each locus were sequenced from a single allotetraploid. *A*: A phasing model which swaps the gene copies between two mul-tree tips (the green and the blue tips) assumes the two gene copies of each locus are homeologous. *B*: An alternative phasing model swaps gene copies between three mul-tree tips. This potentially allows for the gene copies of some loci to be inferred as allelic variation and the gene copies of other loci to be inferred as homeologs. In this example, the gene copies of locus 1 and locus 4 are homeologs and the gene copies of locus 2 and 3 are allelic variants. The fit of the data to these two models—mul-tree A and mul-tree B—can be compared through Bayes factors.

### Software availability

`RevBayes` (Höhnaet al., 2016) is an open-source software package available at http://revbayes.com. RevGadgets (Tribble et al., 2020), the `R` package used for summarizing the output from `RevBayes` analyses, is also open-source and is available at https://github.com/revbayes/revgadgets. Further details, including an example analysis and all scripts and data used in this paper, are available at https://github.com/wf8/homologizer.

### Simulation tests

To evaluate the performance of `homologizer`, we performed two separate simulation tests. The first tested the impact of different evolutionary factors on phasing accuracy and the second focused on whether applying Bayes factors to compare `homologizer` phasing models can successfully distinguish between allelic versus homeologous variation.

#### Experiment 1: Testing phasing accuracy

To explore the behavior of `homologizer` under different evolutionary scenarios, we simulated datasets to test how the performance of `homologizer` was impacted by the following factors: (1) the phylogenetic informativeness of each locus; (2) the phylogenetic distance between subgenomes for each polyploid (the minimum distance in cases of more than two subgenomes); (3) the minimum phylogenetic distance from a polyploid subgenome to the nearest diploid, and; (4) the amount of incomplete lineage sorting (ILS) present. For each replicate we first simulated a species tree under a constant rate birth-death process with root age = 50.0, speciation rate = 0.2, extinction rate = 0.01, and the fraction of taxa sampled at the present = 0.01. Each species tree was conditioned on having twenty extant tips, five of which were randomly selected to represent allopolyploid subgenomes. Two of the allopolyploid tips were then randomly selected to represent the subgenomes of a tetraploid, and the other three allopolyploid tips represented subgenomes of a hexaploid (making the species tree a mul-tree). The remaining 15 tips were considered diploid lineages that did not need phasing. Each dataset thus included two allopolyploids that we could use to assess the performance of `homologizer`.

Once the species tree was simulated, we used the multi-species coalescent to simulate four gene trees, representing four loci. We varied the effective population size of these simulations as described below. Finally, over the simulated gene trees we simulated nucleotide sequences under an independent GTR substitution model with exchangeability rates and stationary frequencies sampled from flat Dirichlet priors. The clock rate of the GTR process was constant over all branches of the tree and fixed to 0.01. We varied the length of the simulated sequences as described below.

All the simulated nucleotide sequences were then analyzed using `homologizer`. The phasing and the phylogeny were inferred simultaneously, linking the tree across loci (so these were “concatenated” analyses). Each locus was modelled by an independent GTR substitution model, with Dirichlet priors on the exchangeability parameters and stationary frequencies, a uniform prior on topology, and exponential priors (mean = 0.1) on branch lengths. The MCMC was run for 2000 generations where each generation consisted of 354 separate Metropolis-Hastings moves. The first 25% of samples were discarded prior to summarizing the posterior distribution. Effective sample sizes (ESS) of the posterior were consistently greater than 200; the mean ESS across simulation replicates was 355.6.

Based on this simulation design, we performed four tests, each varying one of the focal factors while keeping the others at fixed “default” levels. Those default values were a sequence length of 800 base pairs, an effective population size of 0.0001 (which effectively meant there was zero ILS), a minimum distance between polyploid subgenomes of at least 0.25 (scaled by tree height such that a value of 1.0 indicates that two leaves of the tree share a most recent common ancestor at the root of the tree), and a minimum distance from a polyploid subgenome to the nearest diploid of no more than 0.25. For each experiment we tracked the proportion of simulated polyploids for which the phasing was correctly estimated (all four loci were phased correctly), and the mean marginal posterior probability of the joint MAP phasing for each polyploid.

To test the effect of phylogenetic informativeness on the performance of `homologizer`, we varied the length of the simulated nucleotide sequences. For 1000 simulation replicates we sampled the length of the sequence from a uniform distribution with a minimum of 1 and a maximum of 1000 base pairs. To test the effect of the phylogenetic distance between polyploid subgenomes we simulated 800 replicates, without placing any constraints on the maximum subgenome distance. To test the effect of the phylogenetic distance between each polyploid and the closest diploid tip we simulated 800 replicates, this time without constraints on the “diploid distance.” Finally, to test the effect of ILS, we simulated five sets of 200 replicates, with effective population sizes of 0.001, 0.1, 0.5, 1.0, or 2.0, respectively. We then quantified the amount of ILS in each simulated dataset with the summary statistic *R*_*g*_/*R*_*s*_, where *R*_*g*_ is the mean Robinson-Foulds distance (Robinson and Foulds, 1981) between each gene tree and the species tree, and *R*_*s*_ is the mean Robinson-Foulds distance between each independently simulated species tree. This statistic provides an intuitive summary of the amount of gene discordance: when, *e*.*g*., *R*_*g*_/*R*_*s*_ = 0.5 the amount of discordance between the gene trees and the species tree is 50% of what one would observe between completely unlinked trees. Therefore, simulation replicates with 0.0 < *R*_*g*_/*R*_*s*_ ≤ 0.1 represent low amounts of ILS, replicates with 0.1 < *R*_*g*_/*R*_*s*_ ≤ 0.3 represent moderate amounts of ILS, and those with 0.3 < *R*_*g*_/*R*_*s*_ ≤ 0.5 represent high levels of ILS.

#### Experiment 2: Testing Bayes factors to distinguish allelic variation from homeologs

Our second simulation test focused on testing whether Bayes factors could be used to compare `homologizer` phasing models and distinguish allelic variation from homeologs. For each replicate, we first simulated a species tree under a constant-rate birth-death process with root age = 50.0, speciation rate = 0.2, extinction rate = 0.01, and the fraction of taxa sampled at the present = 0.01. Each species tree was conditioned on having twenty extant tips, two of which were randomly selected to represent allopolyploid subgenomes of a tetraploid (making the species tree a mul-tree). The remaining 18 tips were considered diploid lineages that did not need phasing.

Once the species tree was simulated, we used the multi-species coalescent to simulate four gene trees, representing four loci. In half of the replicates, all gene trees were simulated with one gene-tree tip for each species-tree tip. These replicates represent the case in which a two mul-tree tip phasing model is appropriate because there is no allelic variation (all gene copies for the tetraploid are true homeologs). For the other half of the replicates, gene trees were simulated with one gene-tree tip for each species-tree tip for two loci, and with two gene lineages from subgenome A and none from subgenome B for the remaining two loci. These replicates represent the case in which a three mul-tree tip phasing model is needed because the gene copies for two of the loci are allelic variants from the same subgenome but for the other two loci the gene copies are homeologs from different subgenomes.

The gene trees were simulated using an effective population size of 0.0001 (which, like in our first experiment, meant there was zero ILS), a minimum distance between polyploid subgenomes of at least 0.25 (scaled by tree height such that a value of 1.0 indicates two leaves of the tree share a most recent common ancestor at the root of the tree), and a minimum distance from a polyploid subgenome to the nearest diploid of no more than 0.25. Over the simulated gene trees we simulated 800 nucleotide base pairs under an independent GTR substitution model (Tavaré, 1986) with exchangeability rates and stationary frequencies sampled from flat Dirichlet priors. The clock rate of the GTR process was constant over all branches of the tree and fixed to 0.01.

For each simulated dataset we ran stepping-stone analyses to compute marginal likelihoods of the two-mul-tree-tip and three-mul-tree-tip `homologizer` phasing models. Our stepping-stone simulations used 50 stepping stones, sampling 1000 states from each step. We then compared the `homologizer` phasing models using Bayes factors (Kass and Raftery, 1995).

### Cystopteridaceae analysis

Rothfels et al. (2017) analyzed a dataset of four single-copy nuclear loci (*ApPEFP C, gapCpSh, IBR3*, and *pgiC*) for a sample of nine diploids and 19 tetraploids from the fern family Cystopteridaceae, using `alloPPnet` (Jones et al., 2013; Jones, 2017). We reanalyzed these data (Table 1) to compare the performance of `homologizer` with the explicit hybridization model of `alloPPnet`, and to see if the phasing information reported by `homologizer` informs our understanding of this dataset.

**Table 1:**
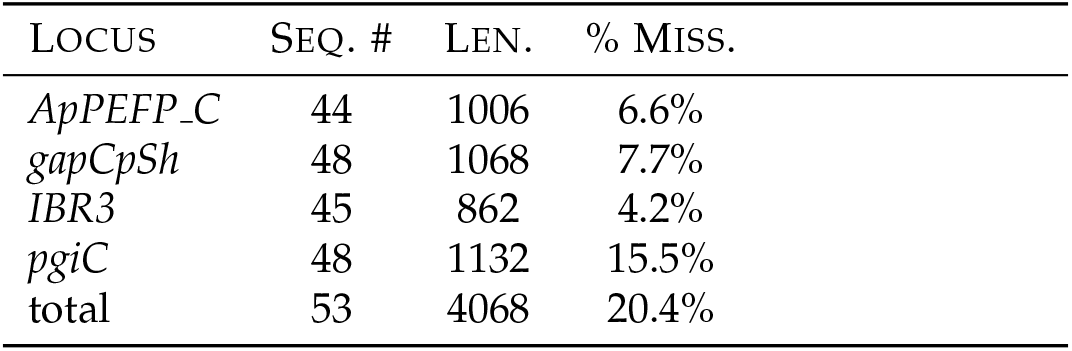
Summary of Cystopteridaceae dataset. SEQ. #: the number of mul-tree tips represented by sequence data for that dataset; LEN.: the aligned length of that dataset in basepairs; % MISS.: the percentage of missing data (?s and -’s).

Because `alloPPnet` cannot accommodate ploidy levels higher than tetraploid, one accession (×*Cystocarpium roskamianum* Fraser-Jenk.; Fraser-Jenkins, 2008; Fraser-Jenkins et al., 2010), which is an intergeneric allotetraploid hybrid of two other allotetraploids and thus includes four distinct evolutionary histories (Rothfels et al., 2015), was treated by Rothfels et al. (2017) as two tetraploids (one for each of its parental genera). With `homologizer`, this ad hoc solution is not necessary, so we reanalyzed these data with ×*Cystocarpium roskamianum* allotted four tips in the mul-tree (to accommodate the homeologs of ×*Cystocarpium*), and most other tetraploids allotted two tips. The diploid samples were each given a single mul-tree tip, except for three (*Cystopteris bulbifera* 7650, *Cystopteris protrusa* 6359, and *Gymnocarpium oyamense* 6399) that had allelic variation and were each treated as two independent accessions. The final dataset comprises 4068 aligned sites (Table 1).

For three of the accessions—*Cystopteris tasmanica* 6379, *Gymnocarpium disjunctum* 7751, and *Gymnocarpium dryopteris* 7981—the number of mul-tree tips required was uncertain (for more discussion of this uncertainty, see the “Insights from `homologizer` model selection and data exploration” section, below). To determine the appropriate number of tips for these taxa, we tested seven different phasing models (Table 2). Two of the models tested whether *C. tasmanica* should have two or three mul-tree tips, another two tested whether *G. disjunctum* should have two or three mul-tree tips, and, finally, three tested whether *G. dryopteris* should have three, four, or five mul-tree tips. We then calculated Bayes factors by stepping-stone analyses (Xie et al., 2011) to compare among the models for each of the three polyploids. We calculated the marginal log-likelihood of each model four times independently; each analysis comprised 50 steps, sampled 1000 time each.

**Table 2:** Cystopteridaceae phasing model tests. Bayes factors were used to compare phasing models for three polyploid accessions. Each model varied the number of mul-tree tips used to phase gene copies. The marginal log-likelihood was calculated four times independently for each phasing model; the mean and standard deviation is shown. The Bayes factor of the best supported model versus each other model is shown. The best supported model is indicated with -. Model comparisons with “strong” support according to Kass and Raftery (1995) are marked with **.

| ACCESSION | NUMBER<br>OF MUL-TREE TIPS | MARGINAL<br>LOG-LIKELIHOOD (MEAN) | MARGINAL<br>LOG-LIKELIHOOD (SD) | BAYES FACTOR |
| --- | --- | --- | --- | --- |
| *C. tasmanica* 6379 | 2 | -14519.12 | 2.27 | **62.34\*\*** |
|  | 3 | -14456.78 | 5.71 | - |
| *G. disjunctum* 7751 | 2 | -14157.80 | 1.81 | 1.90 |
|  | 3 | -14155.91 | 1.82 | - |
| *G. dryopteris* 7981 | 3 | -14216.27 | 2.11 | **26.67\*\*** |
|  | 4 | -14202.02 | 3.02 | **12.42\*\*** |
|  | 5 | -14189.61 | 2.80 | - |

We ensured that each model comparison tested a single polyploid accession in isolation by removing the other two uncertain accessions (i.e., when testing whether *C. tasmanica* should have two or three tips we removed all *G. disjunctum* and *G. dryopteris* tips). Additionally, we fixed the phasing of the other polyploid accessions to the MAP phase previously estimated in an analysis that excluded all three of the focal taxa (results not shown).

After the model comparisons were complete, we chose the number of mul-tree tips with the highest support for each polyploid and then jointly estimated the tree and the phase of all accessions together, linking the tree across loci. Each locus was modelled with an independent GTR substitution model, with Dirichlet priors on the exchangeability parameters and stationary frequencies, a uniform prior on topology, and exponential priors (mean = 0.1) on branch lengths. The MCMC was run seven times independently, each for 10000 generations where each generation consisted of 532 separate Metropolis-Hastings moves. Of these seven runs, three converged (see the “MCMC behaviour for `homologizer` models on empirical data” section, below); one of these runs was arbitrarily selected as our final analysis. The first 10% of samples were discarded prior to summarizing the posterior distribution. The ESS of the posterior was 252. The MCMC log is available at https://github.com/wf8/homologizer/blob/main/data/cystopteridaceae.log.gz.

## Results

### Simulation test results

#### Experiment 1: Testing the phasing accuracy under simulated conditions

`homologizer` was able to correctly phase all four loci, with high posterior probability, over most of our simulation conditions, and in those cases where the joint MAP phasing was incorrect, the posterior probability was generally low (Figure 3). Even with minimal phylogenetic information available (sequence lengths less than 200 base pairs long with the nucleotide substitution process fixed to a rate of 0.01), `homologizer` was able to phase the loci correctly over three-quarters of the time. The only cases where `homologizer` struggled to get the correct phasing was in situations where either the distance between subgenomes was very small, or the distance from the polyploid subgenomes to the nearest diploid was very large (Figure 3, middle two rows). In both those cases, the polyploid subgenomes are likely to be sister to each other and thus there is no topological information available to inform the phasing—the average marginal posterior probability in these cases converges on 50%, as expected (the phasing of each locus in the case of sister lineages being an effectively random choice; Figure 3).

**Figure 3:**
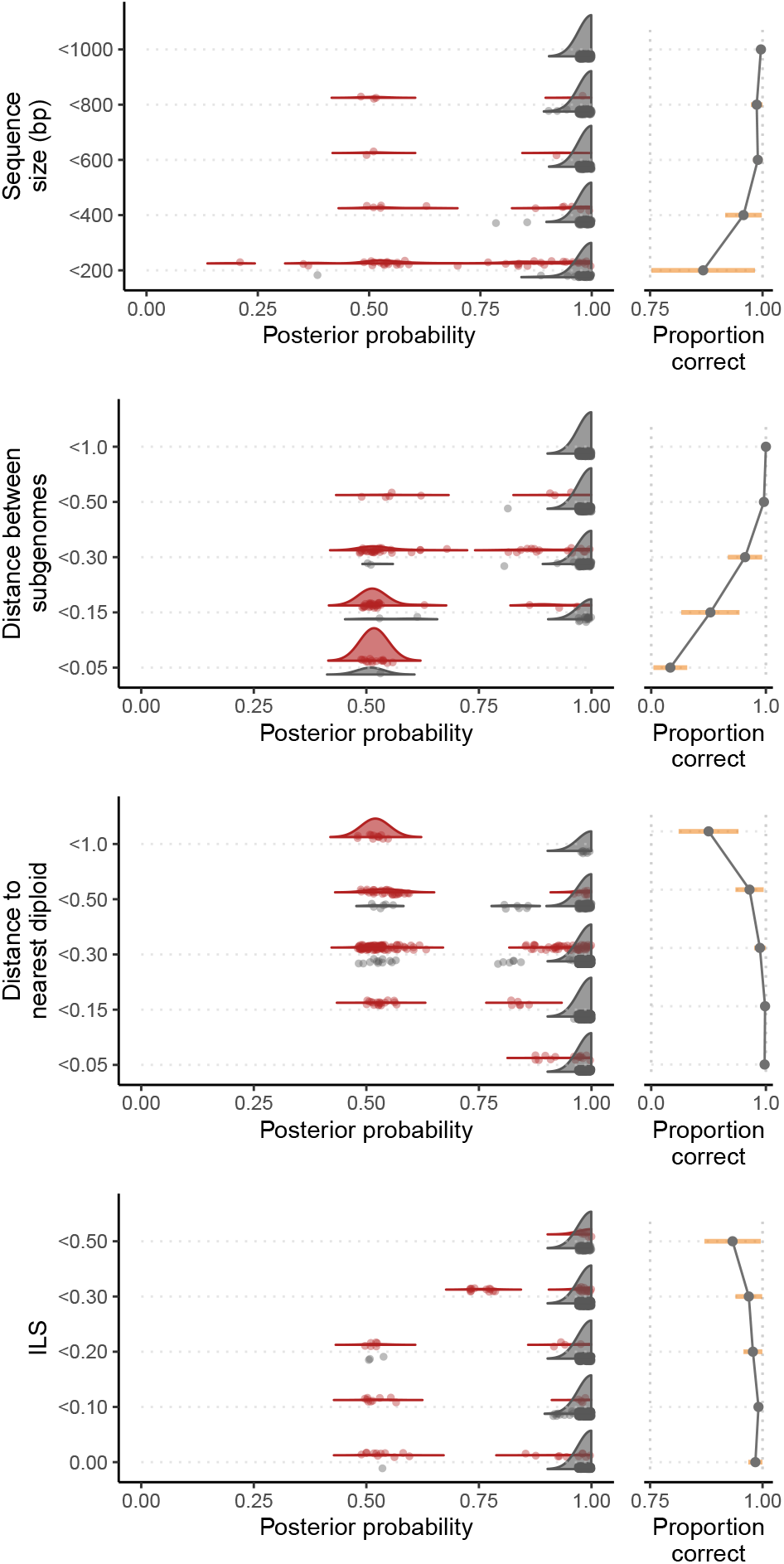
Performance of homologizer in phasing four loci under different simulated conditions. Each row shows how phasing performance was impacted by a factor expected to influence accuracy: sequence length; smallest distance between subgenomes; smallest distance between a polyploid subgenome and a sampled diploid; and degree of incomplete lineage sorting (ILS). See the main text for details on how each factor was quantified. The x-axis in the left panel of each row shows the mean marginal probability of the joint MAP phasing: each point represents an individual simulated polyploid. Points are colored red if the joint MAP phasing was estimated incorrectly and grey if the phasing was estimated correctly. The grey and red densities represent the distribution of the mean marginal posterior probabilities of correctly and incorrectly phased simulated polyploids, respectively. The right panel of each row summarizes how the proportion of times the model was correct changes with the focal factor. The orange bar is the variance.

Gene-tree incongruence (induced in our simulations by elevated degrees of ILS) had a subtly different impact on `homologizer` inference. While the method was generally robust to ILS (Figure 3, bottom row), high levels of incongruence resulted in an appreciable number of simulations where `homologizer` was “confidently wrong”—the joint MAP phasing was incorrect, but the average marginal phasing probability was high. This behavior is due to the model’s assumptions being violated by the process that actually generated the data; the loci evolved along highly discordant gene trees and yet the `homologizer` model assumed a single shared topology for all gene trees. Stochastic variation in the “true” gene trees may result in one phasing having a slight advantage and thus high posterior probability, analogous to the high support inferred for arbitrary resolutions of polytomies by MCMC analyses that only visit fully resolved trees (the “star-tree paradox”; Lewis et al., 2005; Yang, 2007; Rothfels et al., 2012).

#### Experiment 2: Testing Bayes factors to distinguish allelic variation from homeologs

Using Bayes factors to compare `homologizer` phasing models successfully distinguished allelic variation from homeologs (Figure 4). For the simulation replicates in which the three-mul-tree-tip phasing model was necessary to accommodate the presence of both homeologous gene copies and allelic variation, the Bayes factors for the three-multree-tip phasing model compared to the two-mul-tree-tip phasing model were nearly always over 100, indicating “decisive” support (Figure 4, grey dots; Kass and Raftery, 1995). When all gene copies from the tetraploid were homeologs such that three mul-tree tips were not necessary, then the Bayes factors were always less than zero (Figure 4, red dots), indicating there was no support for the three-mul-tree-tip phasing model over the two-mul-tree-tip phasing model.

**Figure 4:**
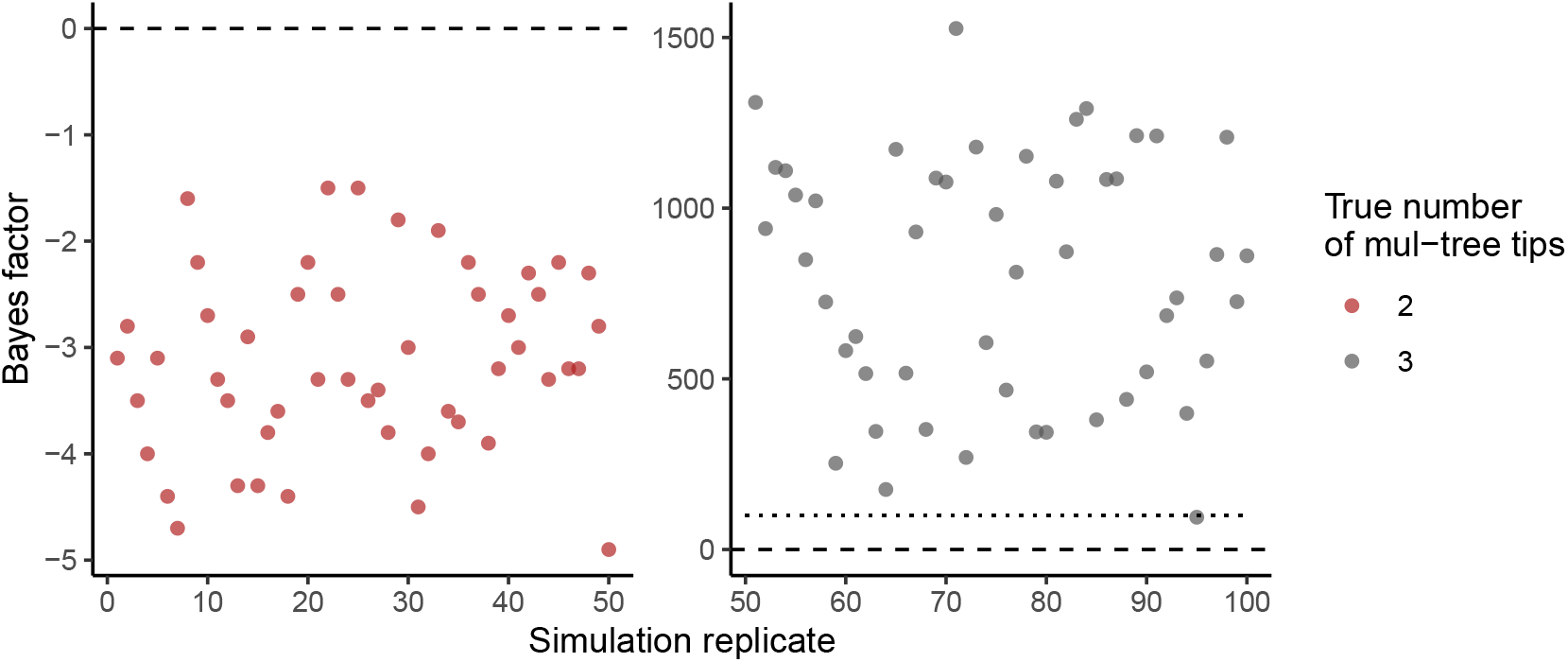
Effectiveness of model comparison tests to distinguish allelic variation from homeologs. Each dot represents a single simulated dataset as described in the main text. The y-axis shows the Bayes factor for the three mul-tree tip phasing model compared to the two mul-tree tip phasing model. All replicates had two gene copies simulated for each locus. Replicates 1–50 (left panel; red dots) were simulated with each of the two tips representing a different polyploid subgenome. The Bayes factor for all of these replicates was less than 0 (dashed line), indicating the three-tip model was not supported over the two-tip phasing model. Replicates 51–100 (right panel; grey dots) were simulated with three total tips representing two different subgenomes of a polyploid in some loci and a single subgenome with two allelic variants in other loci. For nearly all these replicates the Bayes factor was over 100 (dotted line), indicating “decisive” support (Kass and Raftery, 1995) for the three-tip phasing model over the two-tip model.

### Cystopteridaceae analysis results

#### Insights from homologizer model selection and data exploration

The ability to compare among different `homologizer` models with Bayes factors, and thus to use `homologizer` as a data-exploration tool, resulted in key insights about the data for three of the accessions in our dataset; these insights subsequently resulted in significantly altered results of the downstream inferences. The first case involves *Cystopteris tasmanica*, a tetraploid (Tindale and Roy, 2002; Brownlie, 1958; Brownsey and Perrie, 2018). From our *C. tasmanica* accession, we recovered two sequences for three of the four loci (we recovered a single copy from *IBR3*). This accession thus appears to be perfectly “well-behaved”, with one copy of each of the two expected homeologs at most loci, which is how it was treated in the Rothfels et al. (2017) `alloPPnet` analyses. However, preliminary `homologizer` analyses phasing two mul-tree tips for *C. tasmanica* had an odd result: three of the loci phased with very high posterior support, but the phasing for the two *gapCpSh* copies was equivocal (each copy had ∼50% posterior probability of phasing to each of the mul-tree tips). To test the possibility that this strange result was due to the two *gapCpSh* copies being allelic rather than homoelogous (in which case we would expect each of the copies to fit equally to each mul-tree tip), we ran another `homologizer` model, this time with *C. tasmanica* allocated three mul-tree tips. This three-tip model strongly out-performed the two-tip one (Table 2), and as we had suspected, the two *gapCpSh* copies were resolved sister to each other, with a blank copy phased with high posterior probability to the other subgenome (Figure 5).

**Figure 5:**
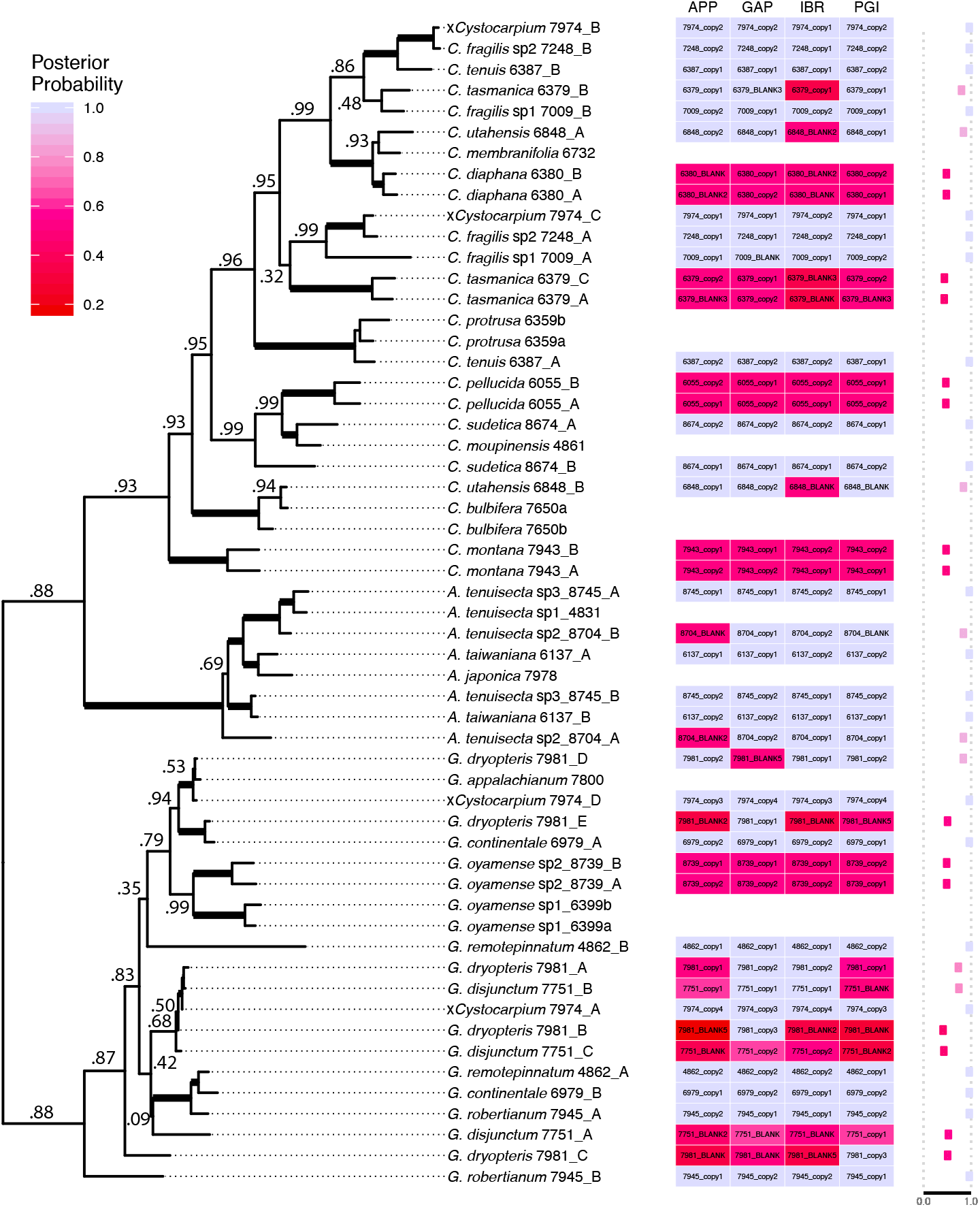
homologizer analysis of the Cystopteridaceae dataset. The phasing of gene copies into subgenomes is summarized on the maximum *a posteriori* (MAP) phylogeny. Thickened branches have posterior probabilities of 1.0; posterior probabilities < 1.0 are indicated. The columns of the heatmap each represent a locus, and the joint MAP phase assignment is shown as text within each box. Each box is colored by the marginal posterior probability of the phase assignment. Adjacent to the heatmap is a column that shows the mean marginal probability across loci of the phasing assignment per tip, which summarizes the model’s overall confidence in the phasing of that tip. In the sample labels, “A.” = *Acystopteris*, “C.” = *Cystopteris*, “G.” = *Gymnocarpium*, the four-digit numbers are Fern* Labs Database accession numbers (https://fernlab.biology.duke.edu/), capital A, B, etc, indicate subgenomes, sp1 and sp2 indicate undescribed cryptic species, and lowercase letters following the accession numbers indicate haploid “individuals” within the sampled diploids (*i*.*e*., those diploids are heterozygous). Copy names with a “BLANK” suffix indicate missing sequences (*e*.*g*., a subgenome that was present in some loci but not retrieved for others).

The other two insights were related to each other, and involve the allotetraploid *Gymnocarpium dryopteris* and one of its diploid progenitors, *G. disjunctum* (Pryer and Haufler, *1993; Rothfels et al*., *2014, 2017). In our dataset, G. dryopteris* was represented by two sequences for two loci (as would generally be expected for an allotetraploid), but by three sequences for the other two loci. In the `alloPPnet` multispecies coalescent analyses of Rothfels et al. (2017), this accession was split into two “individuals” in order to accommodate its apparent heterozygosity. For our preliminary concatenation-based (instead of coalescent-based, as in `alloPPnet`) `homologizer` analyses, we ran a model that gave this accession three tips, in the thought that only one of the subgenomes had any allelic variation (*i*.*e*., the pattern shown in Figure 2B), and a four-tip model, in case both subgenomes were heterozygous at at least one locus. The phasing for *G. dryopteris* in these preliminary analyses, however, had difficulty converging, and in addition, inspection of the *pgiC* gene tree showed that each of the three *G. dryopteris* copies were resolved in distinct places. Specifically, one copy grouped with each of *G. appalachianum* and *G. disjunctum*, as expected given these two diploids are the *G. dryopteris* progenitors (Pryer and Haufler, 1993; Rothfels et al., 2014, 2017); however, the third copy was in an isolated position. In response, we ran a five-tip model, and this model was strongly preferred over either of the three- or four-tip ones (Table 2). Similarly, *G. disjunctum* appeared to be heterozygous at two loci and in Rothfels et al. (2017) was split into two individuals; however this (two-tip) model converged poorly, and a three-tip model was weakly preferred by Bayes factor comparison (Table 2).

The final, integrated model (three tips for *C. tasmanica*, five tips for *G. dryopteris*, and three for *G. disjunctum*) resulted in a considerably revised understanding of our dataset. The *C. tasmanica* case was fairly straightforward, but also had a dramatic effect: by allowing the *gapCpSh* copies to be allelic rather than forcing them onto different subgenomes, the three-tip model inferred that *C. tasmanica* is an allopolyploid, with one subgenome from each of the major subclades of the core *fragilis* complex (*i*.*e*., the *fragilis* complex minus *C. protrusa*; Rothfels et al., 2013). This conclusion is in strong contrast to that of Rothfels et al. (2017), which limited this accession to two tips and inferred that it was possibly an autopolyploid (the two tips were inferred to be sister to each other, albeit relatively deeply divergent). For *G. disjunctum*, both sequences of the two loci that had two copies (*gapCpSh, IBR3*) were phased to tips consistent with their being “normal” allele variants in a diploid genome—these two loci are heterozygous. However, the lone *pgiC* sequence did not phase to either of those tips; instead it was phased to its own tip, in an isolated portion of the phylogeny (Figure 5). Multiple lines of evidence suggest that this tip does not represent a subgenome history: 1) it is inconsistent with what is known of the ploidy of *G. disjunctum* (it’s a diploid; Pryer and Haufler, 1993; Rothfels et al., 2014); 2) it is inconsistent with the phylogenetic history of this taxon (as inferred from multiple accessions of *G. disjunctum* itself as well as from the allopolyploids of which it is a genome donor (×*Cystocarpium* and *G. dryopteris*; Rothfels et al., 2014, 2017)), and; 3) this history is not represented by any of the other loci (Figure 5). Based on that evidence, and particularly in the context of the results for *G. dryopteris* (below), we suspect that this sequence represents a lineage-specific duplication of *pgiC* within the *G. disjunctum* lineage.

For *G. dryopteris*, while each locus had only one sequence from the subgenome closely related to *G. appalachianum*, the *gapCpSh* representative of this subgenome was sufficiently phylogenetically distinct (presumably due to coalescent variance) to be phased with high posterior probability to its own, closely related, tip. In the subgenome closely related to *G. disjunctum*, the results were more to be expected—all loci had a single copy, which phased to one of the tips, except for *gapCpSh*, which had two closely related alleles (Figure 5). However, the most important result was that one of the three *pgiC* copies was phased, with high posterior probability, to its own tip, phylogenetically distant from all other *G. dryopteris* tips. This pattern is very similar to that seen for *pgiC* in *G. disjunctum* (above), and we suspect these issues are related: *G. dryopteris* has inherited this putatively paralogous copy from *G. disjunctum*, and the *G. dryopteris* version in our dataset may additionally be a PCR recombinant. This conclusion is supported by the fact that *G. dryopteris* has two other *pgiC* copies that are cleanly phased to subgenomes corresponding to each of the putative progenitor diploids. By allowing this aberrant *pgiC* tip to be phased to its own branch, this three-tip `homologizer` model clarifies that the remaining *G. disjunctum* allele sequences are very closely related to each other, and provides strong support to the relationships within this area of the *Gymnocarpium* phylogeny (Figure 5).

#### MCMC behaviour for homologizer models on empirical data

The Cystopteridaceae analyses also demonstrated that inference under `homologizer` models, when applied to real datasets with a large number of polyploid accessions, involves complex interactions among the phasing estimates and between those estimates and those of the topology (and branch lengths). Particularly problematic is the case of two polyploids that have closely related subgenomes and can get their phasing for a subset of the loci “stuck” in the incorrect orientation. In the Cystopteridaceae dataset, for example, ×*Cystocarpium* and *Cystopteris fragilis* “sp 2” share two pairs of closely related subgenomes, each pair from a different major subclade of the *fragilis* complex (Figure 5). If a locus got out-of-phase for both accessions, it could get trapped in that orientation, presumably because a proposal to the phasing for that locus for any one of the accessions would result in each pair of closely related tips containing very dissimilar sequences, which would have a low probability given the short phylogenetic distance between the two accessions. While we experimented with more complex joint phase moves, we found that simply increasing the frequency of the phase moves relative to the tree topology moves was more effective. However, while any given `homologizer` run was reasonably rapid (on the order of a “regular” analysis in `RevBayes`), they frequently failed to converge. Specifically, individual runs would get stuck in a suboptimal region of parameter space and appear to be sampling from the posterior distribution (effective samples sizes would be high, etc.), but multiple such runs would sample from different distributions. In our full Cystopteridaceae analysis, for example, we ran seven independent runs, only three of which converged to the posterior distribution. As a result, we recommend that users run preliminary analyses and inspect the results for areas where two closely related accessions are potentially interfering with each other. In addition, multiple independent runs are necessary to ascertain the efficiency of MCMC mixing for a given dataset. Users can additionally experiment with Metropolis-coupled Markov chain Monte Carlo (as implemented in `RevBayes`; Altekar et al., 2004) to improve mixing.

#### A phased multi-locus Cystopteridaceae phylogeny

Our inferred Cystopteridaceae phylogeny under the best-fitting model (see Table 2) is well supported (Figure 5), and generally consistent with the inferences of Rothfels et al. (2017), although with the addition of all the phasing information.

The primary differences relate to the insights gained from our data exploration/model comparison (see above). Specifically, the strongly supported resolution of *Cystopteris tasmanica* as an allopolyploid with two fairly phylogenetically distant progenitors (Figure 5) is in strong contrast to the earlier inferences of this accession as a potential autopolyploid, with subgenomes that are sister to each other (Rothfels et al., 2017). Other secondary differences include the resolution of the *sudetica* clade sister to the *fragilis* complex instead of to the *bulbifera* clade, and a different placement of the morphologically anomalous *G. oyamense* within *Gymnocarpium*. It is difficult to determine what is driving these (secondary) topological differences, since, beyond the phasing component, the underlying analyses differ in their substitution model, clock model (or lack thereof), and in whether or not they model ILS (through the multi-species coalescent). Furthermore, while these relationships are all strongly supported in our `homologizer` analysis, `alloPPnet` doesn’t report clade probabilities, so it is not clear how well supported the alternative relationships were in the earlier `alloPPnet`-based analysis.

The phasing is also generally well supported, despite the fact that considerable portions of the phylogeny don’t have any diploid representation: by sharing phylogenetic information among polyploids we were able to phase them confidently even without closely related diploids being included. Those subgenome pairs that phased poorly (had low average marginal probabilities of phasing) were restricted to sister-pairs, where there is no topological phasing information available. Low marginal probabilities of phasing for individual loci (*e*.*g*., where three loci are phased with high probability but the forth is not) nearly always reflect cases where that locus was not recovered for the given sample, such as IBR for *C. utahensis* (Figure 5).

## Discussion

### homologizer for inference

The core application of `homologizer` is to infer multilocus multilabeled phylogenies and/or the phasing of gene copies to subgenomes. For this application, our simulation tests show that `homologizer` is robust to a wide range of factors that can potentially affect accuracy (Figure 3). When the `homologizer` model does not have enough information to confidently phase gene copies due to (1) phylogenetically uninformative sequences, (2) the polyploid subgenomes being too closely related, or (3) a lack of diploid lineages that help inform the phasing, then `homologizer` makes estimates with low posterior probability that correctly capture the uncertainty of the phasing.

Model violations, however, pose a greater challenge to `homologizer`. We see this sensitivity in our simulations with high levels of ILS (Figure 3, bottom panel). Here, since the `homologizer` model used in our simulation tests assumed a shared gene tree, the incongruence among gene trees was not modeled appropriately, and `homologizer` sometimes made incorrect estimates with high posterior probability (“confidently wrong” behavior). However, the modularity of `RevBayes` allows for the easy integration of `homologizer` with other model components. For example, if a user expects ILS to be important in their dataset, we recommend using `homologizer` within the full multi-species coalescent (MSC) framework, allowing for unlinked gene trees. This model, however, is computationally demanding (like all full MSC models), and can feasibly be applied to only relatively small datasets. Another type of model violation that may mislead homologizer analyses is recombination among the subgenomes within a polyploid. Recombination could potentially result in chimeric gene copies consisting of fused together portions of different subgenomes, which would violate the assumption of a bifurcating evolution history for the recombined locus. However, recombination among the subgenomes in relatively recently formed polyploids (i.e, “neopolyploids”) is uncommon, such that most polyploids are characterized by “fixed heterozygosity” (Sigel, 2016); recombination within a focal locus would be very rare (Chen et al., 2018).

Other datasets might require different analysis approaches. For example, genome-scale target enrichment and transcriptome datasets often contain hundreds of loci. With these large datasets joint inference of the phylogeny and gene-copy phasing as presented here may not be computationally feasible. In those cases, it may be reasonable to adopt a sequential Bayesian approach: first infer a species-level mul-tree while phasing a subsample of loci, and then condition the phasing of the remaining loci on that mul-tree. During the secondary phasing analyses one could optionally incorporate phylogenetic uncertainty in the mul-tree by integrating over the posterior distribution of mul-trees, as inferred in the first step. The `RevBayes` implementation of `homologizer` allows for a wide range of approaches, such as these, for scaling up phasing analyses that should be suitable for any sized dataset.

### homologizer for data exploration

The other core functionality of `homologizer` is as a data exploration tool. With increasing numbers of taxa and loci, it can be difficult to detect non-orthologous sequences, such as hidden paralogs, contaminants, or allelic variation that is erroneously modelled as homeologous. `homologizer`, in conjunction with the `RevGadgets` visualization tools, allows for the convenient detection of cases where most loci phase easily but some do not, indicating a likely problem in the homology of the underlying sequences. This application can be extended as a hypothesis-testing tool via model comparison (see “Experiment 2: Testing Bayes factors to distinguish allelic variation from homeologs”), which provides an objective, powerful, and statistically rigorous method of understanding these complex patterns of homology. In the case of our empirical Cystopteridaceae dataset, this data exploration approach revealed cases of alleles masquerading as homeologs (*e*.*g*., in *Cystopteris tasmanica*) and of hidden paralogy (in *Gymnocarpium disjunctum* and *dryopteris*). The identification and correction of these errors resulted in important changes to our downstream phylogenetic inference, demonstrating the impact that this form of data exploration can have.

## Summary

`homologizer` provides a powerful and flexible method of phasing gene copies into subgenomes, allowing users to infer a multilocus phylogeny for lineages with polyploids (or F1 hybrids) while treating the phasing as a nuisance parameter, to infer the phasing while integrating out the phylogeny, or for any combination of these goals. Our simulations demonstrate that `homologizer` is able to confidently infer phasing even in cases of relatively weak phylogenetic signal (*e*.*g*., short loci); in such cases where the signal is insufficient, `homologizer` returns an equivocal result (all potential phasings have lower posterior probability). In contrast, our simulations also demonstrate that `homologizer` can be sensitive to model violations. Specifically, if the true gene trees differ strongly from each other (*e*.*g*., due to high levels of ILS) and yet the model assumes all loci share a single phylogeny, the method is forced to choose among a small set of more-or-less equally wrong phasing options, and may return the incorrect phasing with a high posterior probability (*i*.*e*., the best of a bad lot). Our analyses of the empirical Cystopteridaceae dataset further demonstrate the power of the method; by simultaneously leveraging the phylogenetic information available in the full taxon sample (diploids and polyploids alike), across all loci, `homologizer` can confidently infer the phylogeny and phasing in areas of the tree where there is no diploid representation, and for markers where there is extensive missing data. As such, `homologizer` opens up the application of multilocus phylogenetics to groups containing polyploids or other individual organisms that may contain subgenomes with divergent evolutionary histories—such groups have been historically understudied phylogenetically, which has limited our ability to explore evolutionary questions in general, and questions related to the macroevolution of polyploids specifically. Finally, `homologizer` may potentially be used beyond its core phasing functionality to identify non-homologous sequences more generally, with broad applications for phylogenetic and phylogenomic inference.

## Acknowledgements

The authors would like to thank Bruce Baldwin for helpful discussions that inspired this work, and the Rothfels lab for comments that significantly improved the manuscript.

## Supplementary information

A full tutorial for setting up and running `homologizer` analyses is available at https://github.com/wf8/homologizer, but we provide some details here.

### Setting up a homologizer analysis

Two types of information are required in order to run a `homologizer` analysis. First are the sequence data—the multiple-sequence alignments for each of the loci. Note that the sequences within these alignments should already have been correctly assembled into haplotypes using standard phasing approaches. Second is a determination, by the user, of how many tips each accession should get in the mul-tree. For example, if an accession has more than one copy at one or more loci, the user needs to decide whether those copies potentially represent multiple subgenomes or whether they instead represent allelic variation (in which case the user might split them into separate “individuals”— separate distinct tips that are not available for gene-copy phasing). These determinations (the number of tips among which to phase gene copies) are fixed for a given analysis, although the tools of Bayesian model selection can be applied to determine how many tips are justified for each accession (see Model Selection section, below).

`homologizer` is implemented in `RevBayes` (Höhnaet al., 2016), which is an modular program for implementing Bayesian phylogenetic analyses that uses an `R`-like syntax (R Core Team, 2013). Included in `RevBayes`’ core functionality are tools for reading multiple sequence alignments and setting up all elements of a phylogenetic analysis (the substitution models, tree models, clock models, etc). For example, for the Cystopteridaceae analyses included in this paper, we set up an independent GTR substitution model for each locus, and ran a concatenated analysis (a single mul-tree topology shared by all loci). For the `homologizer` component of the model, a user initializes the MCMC analysis by first seting an initial phase assignment for each of the copies to be phased, using the `setHomeologPhase` command. For example, the Cystopteridaceae dataset has an accession, ×*Cystocarpium* #7974, that has four sequences for locus1, labelled “7974 copy1” through “7974 copy4”, and we wish to phase those copies among four mul-tree tips, labelled “xCystocarpium 7974 A” through “xCystocarpium 7974 D”. To set the initial phase assignment for these copies, one would use the following commands:

locus1 . setHomeolog Phase (“7974 _copy1”, “x Cystocarpium_7974 _A “)

locus1 . setHomeolog Phase (“7974 _copy2”, “x Cystocarpium_7974 _B “)

locus1 . setHomeolog Phase (“7974 _copy3”, “x Cystocarpium_7974 _C “)

locus1 . setHomeolog Phase (“7974 _copy4”, “x Cystocarpium_7974 _D “)

To successfully initialize the MCMC, each gene copy needs to have a unique initial phase assignment (each copy needs to be assigned to a mul-tree tip, and no two copies may be assigned to the same tip). This initial phase assignment can be randomly picked by the user and is only used to initialize the MCMC; it will not affect the final phasing estimate assuming the MCMC analysis converges.

The next command needed to set up a `homologizer` analysis is `mvHomeologPhase`, which creates an MCMC proposal that tries swapping gene copies among the mul-tree tips of a polyploid. For each locus with *N* gene copies to be phased, the analysis requires 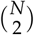 MCMC phase swapping proposals, so for ×*Cystocarpium*, with its four mul-tree tips, we need to apply the `mvHomeologPhase` command six times for each locus:

moves [++ mvi] = mv Homeolog Phase (locus1, “x Cystocarpium_7974 _A”, “x Cystocarpium_7974 _B “)

moves [++ mvi] = mv Homeolog Phase (locus1, “x Cystocarpium_7974 _A”, “x Cystocarpium_7974 _C “)

moves [++ mvi] = mv Homeolog Phase (locus1, “x Cystocarpium_7974 _B”, “x Cystocarpium_7974 _C “)

moves [++ mvi] = mv Homeolog Phase (locus1, “x Cystocarpium_7974 _A”, “x Cystocarpium_7974 _D “)

moves [++ mvi] = mv Homeolog Phase (locus1, “x Cystocarpium_7974 _B”, “x Cystocarpium_7974 _D “)

moves [++ mvi] = mv Homeolog Phase (locus1, “x Cystocarpium_7974 _C”,” x Cystocarpium_7974 _D “)

The gene copy phase assignments sampled during the MCMC are logged to files using a monitor created with the `mnHomeologPhase` command.

There are complex interdependencies among the phasing estimation across accessions that can make it difficult for the MCMC to sample efficiently. For example, two closely related tips can get the phasing for a subset of their loci “stuck”, because switching the phasing to the correct orientation requires traversing a low-probability valley in parameter space where one of the accessions has the correct phasing but the other does not. Users should run multiple independent MCMC analyses and check for convergence.

